# Oxygen level alters energy metabolism in bovine preimplantation embryos

**DOI:** 10.1101/2024.10.20.619258

**Authors:** N. Boskovic, M. Ivask, G. Yazgeldi Gunaydin, B. Yaşar, S. Katayama, A. Salumets, T. Org, A. Kurg, K. Lundin, T. Tuuri, C. O. Daub, J. Kere

## Abstract

**STUDY QUESTION:** What is the effect of different oxygen (O_2_) levels on the transcriptomic profile of bovine embryos during the *in vitro* culture?

**SUMMARY ANSWER:** Embryos grown in hypoxia (6% O_2_) from zygotes until the blastocyst stage had the highest blastocyst formation rate, whereas normoxia (20% O_2_) delayed transcriptomic reprogramming and embryonic genome activation, and induced changes in energy metabolism gene expression.

**WHAT IS KNOWN ALREADY:** Mammalian preimplantation embryo development is a complex sequence of events where, within a week, the zygote is reprogrammed to totipotency and subsequently diverges to embryonic and extraembryonic cell lineages for post-implantation embryo development. This period of development is sensitive to oxygen levels that can affect various cellular processes.

**STUDY DESIGN, SIZE, DURATION:** In this study, we used triplicate bovine embryos as a model for human embryogenesis to compare the influence of O_2_ levels on preimplantation embryonic development by culturing embryos either in normoxic (20% O_2_) or physiological hypoxic (6% O_2_) conditions, or sequential hypoxia until 16-cell stage and then switching to ultrahypoxic culture (2% O_2_).

**PARTICIPANTS/MATERIALS, SETTING, METHODS:** As the readout for varied O_2_ effects, we performed RNA sequencing using 5’ targeted STRT-N method on single embryos. We compared zygotes, 4-, 8-, 16-cell and blastocyst stage embryos grown in either normoxic or hypoxic condition, adding ultrahypoxia for blastocyst stage embryos as the third condition.

**MAIN RESULTS AND THE ROLE OF CHANCE:** We found that the initial cleavage rate was not affected by O_2_ levels but there was a clear difference in blastocyst formation rate. In hypoxia, 36% of embryos reached blastocyst stage while in normoxia the blastocyst formation rate was 13%. In final ultrahypoxia condition only 4.6% of embryos reached blastocyst stage. Transcriptomic profiles showed that normoxic conditions slowed down oocyte transcript degradation and embryonic genome activation. Key metabolic enzyme genes were also altered between hypoxic and normoxic conditions at the blastocyst stage. Both hypoxic and ultrahypoxic conditions induced energy production by upregulating genes involved in glycolysis and lipid metabolism typical to *in vivo* embryos. In contrast, normoxic conditions failed to upregulate glycolysis genes and only depended on primitive oxidative phosphorylation metabolism. We conclude that constant hypoxia culture of *in vitro* embryos provided the highest blastocyst formation rate and appropriate energy metabolism. Normoxia altered the energy metabolism and decreased the blastocyst formation rate. Even though ultrahypoxia at blastocyst stage resulted in a drop of blastocyst formation, the transcriptional profile of surviving embryos was normal.

**LARGE SCALE DATA:** The raw data (BCL files) are available at Zenodo: XXXXX FASTQ files generated are available in the EMBL’s European Bioinformatics Institute (EMBL-EBI)-BioStudies with accession number X-XXXX.

**LIMITATIONS, REASONS FOR CAUTION:** The limitation of this study is the use of bovine as an animal model instead of human embryos. Due to this, the direct translation of the results to human should be taken with caution.

**WIDER IMPLICATIONS OF THE FINDINGS:** This study supports previous literature on hypoxic culture conditions being the most suitable for *in vitro* embryo culture. In addition, we provide new insights on why embryos grown in normoxia do not have the same success rate as embryos grown in hypoxia. We did not observe any benefits of lowering the oxygen levels to 2%, calling for caution of switching to this culture system.

**STUDY FUNDING/COMPETING INTEREST(S):** This project has received funding from the European Union’s Horizon 2020 Research and Innovation Programme under the Marie Sklodowska-Curie grant agreement No. 813707.; Work in the JK laboratory is supported by Jane and Aatos Erkko Foundation, Sigrid Jusélius Foundation, Liv och Hälsa (Finland), Swedish Brain Foundation and Swedish Research Council. The study was also supported by the Estonian Research Council (grant no. PRG1076) and the Horizon Europe NESTOR project (grant no. 101120075).

**What this means for patients?:** During assisted reproduction treatment (ART) oocytes are fertilized in vitro by sperm to form an embryo. The embryos are then exposed to environmental effects during the 5 days of culture before implantation. Embryo culture takes place inside the incubator, where different oxygen levels can be used.

In this study, we compared 3 different oxygen concentrations for embryo culture: atmospheric or normoxia (20%), physiologic or hypoxia (6%) and combined low and ultra-low concentration with initial culture at 6% oxygen until day 3 followed by 2% (ultrahypoxia) until day 5 of culture. We conducted this study because many ART clinics use different oxygen concentrations for embryo culture, and we aimed to provide insight on how these different concentrations impact embryo development by monitoring gene expression.

As using human embryos for this type of experiments is not commonly approved, we used bovine embryos whose early development resembles that of humans. We found that culturing embryos for 5 days at constant low oxygen concentration (hypoxia) gives the highest number of embryos that reach the blastocyst stage (day 5) and have expected gene expression that provides the physiological path of embryo development. Culturing embryos at atmospheric oxygen concentration showed that only 13% of fertilised eggs reached the blastocyst stage at day 5, while culturing at the final ultra-low oxygen concentration had the lowest percentage of matured embryos, and we did not observe any benefits that would justify the use of such a culture system. We suggest that *in vitro* culturing of embryos at constant 5-6% oxygen gives the best results.

## Introduction

Early preimplantation embryo development is a highly coordinated and complex process with a cascade of precisely timed events. Different from the rather stable *in vivo* conditions, *in vitro* embryos are exposed to distinct environmental factors such as culture medium composition, pH, temperature, and oxygen (O_2_) levels. Culture medium and its composition has been shown to affect embryonic metabolism in mouse, rabbit and bovine (Donjacour *et al*., 2014; Sun *et al*., 2015; Wang *et al*., 2022; Pasquariello *et al*., 2023), and it also has an impact on the success rate of human *in vitro* fertilization (IVF) and on the birthweights of infants born after IVF (Kleijkers *et al*., 2016).

The importance of O_2_ concentration during *in vitro* culture (IVC) of embryos was first understood in the first report of successful human embryo culture (Steptoe *et al*., 1971) where oxygen tension was maintained at 5%. In several mammalian species, O_2_ tension in female reproductive tract (oviducts and uterus) ranges generally between 2-8% (Fischer and Bavister, 1993). Despite this knowledge, 49% of clinical IVF laboratories in Europe and 32% in the USA use atmospheric 20% O_2_ levels in their embryo cultures (Christianson *et al*., 2014; Sciorio and Rinaudo, 2023). Recently, many meta-analyses pointed out that clinical pregnancy rates were clearly higher if embryos were cultured in physiological 5% O_2_ (32-43%) rather than in atmospheric 20% O_2_ (30%) (Bontekoe *et al*., 2012).

The epigenetic, metabolic, and transcriptomic profiles of embryos grown under different O_2_ levels have been studied in different species. These studies focused on comparing the low monophasic O_2_ level to the atmospheric O_2_. In the mouse, for example, atmospheric O_2_ changes the number of mitochondria and their structure in the blastocyst stage (Belli *et al*., 2019) as well as the glucose and amino acid metabolism (Kelley and Gardner, 2019). In addition, the impact of atmospheric O_2_ was seen as early as in 2-to 4-cell mouse embryos during the time of embryonic genome activation (EGA) as the inhibition of histone lactylation (Yao *et al*., 2024).

Interestingly, the level of O_2_ is estimated to be the lowest inside the uterus (2% O_2_), motivating the question whether a further decrease in the level of O_2_ after day 3 of development might be more beneficial for the blastocyst formation and pregnancy rates (Morin, 2017). In studies with human embryos focusing on such ‘sequential’ low O_2_ cultures, ultra-low O_2_ culture conditions were not found to be superior to monophasic low O_2_ (Kaser *et al*., 2016; Yang *et al*., 2016; De Munck *et al*., 2019).

To understand the impact of different O_2_ levels on the transcriptome, we cultured embryos in either atmospheric 20% O_2_, hypoxic 6% O_2_ or in the sequential low O_2_ where embryos were first cultured at 6% O_2_ until they reached 16-cell and then moved to 2% O_2_ until the blastocyst stage. We employed the 5’ end-targeted RNA sequencing method, STRT-N, which allows us to distinguish between the degrading maternal and the first upregulated embryonic transcripts and compared zygote, 4-, 8-, 16-cell embryos, grown in either normoxic or hypoxic conditions. Blastocyst stage embryos were compared across 3 conditions: normoxia, hypoxia and sequential ultrahypoxia (2% O_2_).

Considering ethical issues with human embryos, we used bovine embryos as an animal model. The advantages of using bovine model for human development are reviewed in Santos *et al*., (2014). In bovine, oogenesis takes several months while oocyte maturation takes 20-24 h, and the size of the matured oocyte is around 120 μm (Otoi *et al*., 1997). These processes are significantly shorter in the mouse. During human EGA, the first active genes are PRD-like homeobox genes (Töhönen *et al*., 2015). These genes are absent from mouse (Jouhilahti *et al*., 2016), but present in bovine (Yaşar et al., 2024; unpublished data), supporting the usefulness of bovine embryos as a model for human development.

## Materials and methods

### Ovary collection and oocyte extraction

Bovine ovaries were collected in 37°C prewarmed 0.9% sterile NaCl solution and transported from the local slaughterhouse to the laboratory. The ovaries were washed twice with fresh, warm, sterile 0.9% NaCl solution before continuing with the cumulus oocyte complexes (COCs) aspiration from the 2-8 mm size follicles using vacuum pump (Minitüb GmbH). COCs grade 1 (Bó and Mapletoft, 2018) were then cultured in groups of 50 COCs in 500 μl in *in vitro* maturation (IVM) medium containing 0.8% of bovine serum albumin (BSA), 0.5 mM pyruvate, 1 mM of L-glutamine, 0.05 μg/ml of EGF in 4-well Nunc plates (Thermo Fisher Scientific). Based on the condition tested, IVM was performed for 22-24 h at 38.5°C, 5% CO_2_.

### *In vitro* fertilization (IVF)

The matured oocytes were fertilized with thawed frozen semen (Holstein breed bull) by co-incubating them in groups of 50 in 500 μl IVF medium supplemented with 0.6% BSA, 0.25 mM pyruvate, 20 μM penicillamine, 10 μM hypotaurine, 1 μM epinephrine and 30 μg/ml heparin. The 4-well Nunc plates were placed at 38.5°C, 5% CO_2_ for 19-20 h.

### Embryo culture and collection

After IVF, cumulus cells were removed by vortexing, denuded zygotes were transferred to 500 μl of modified Synthetic Oviduct Fluid (SOF) supplemented with 0.8% BSA, 0.25 mM pyruvate, 0.2 mM L-glutamine, BME (Sigma-Aldrich, B6766) and MEM amino acid solutions (Sigma-Aldrich, M7145). *In vitro* culture (IVC) was performed based on the testing conditions. Two 4-well Nunc plates were placed in the incubator at 20% O_2_ for 8 days in normoxia condition. For hypoxia, embryos were cultured at 6% O_2_ the whole 8 days, while for the ultrahypoxia condition embryos were cultured until 16-cell stage at 6% O_2_ and then placed to 2% O_2_ until the day 8 of development.

One 4-well control dish per condition, used to calculate cleavage rate and blastocyst formation rate, was left in the incubators for 8 days and only removed at the time of checking the cleavage and blastocyst formation rates (Supplementary table 2). The rates were calculated based on each well in the control dish using the number of presumptive zygotes per well. The other plates were used to collect embryos for RNA sequencing. Three embryos per developmental stage (zygote, 4-cell, 8-cell, 16-cell, and blastocyst) were collected and placed in the cell lysis buffer as previously described in Boskovic *et al*., (2023). Briefly, embryos were taken out of the IVC plate and placed in Acid Tyrode’s solution until *zona pellucida* was dissolved. Then embryos were washed in a drop of IVC medium, followed by washing in two consecutive drops of PBS before placing it in the cell lysis buffer.

### Single oocyte/embryo RNA-seq library preparation

Single embryo RNA-seq libraries were prepared using the STRT-N seq protocol (Boskovic *et al*., 2023). In brief, cell lysate plate was removed from -80°C and RNA was immediately denatured for 2 min at 80°C and placed on ice. Reverse transcription mixture was added containing synthetic ERCC Spike-ins mix. Reverse transcription was performed at 42°C for 1 h followed by 10 cycles of 42°C and 50°C each lasting 2 min. cDNA was cleaned up and barcoded primers were added before performing cDNA amplification with 23 PCR cycles. cDNA samples were pooled, cleaned up and fragmented. Samples were then prepared for sequencing by extracting the 5’ end of the cDNA, repairing the ends and ligating the sequencing adapters, before performing the final library amplification.

### Processing and analysis of the STRT-N RNA-seq libraries

Preprocessing of the raw RNA-seq data was done on Center for Scientific Computing (CSC) HPC server (Finland) using the pipeline described in https://github.com/gyazgeldi/STRTN, (commit number 5b60c80) (Boskovic *et al*., 2023). Raw reads were mapped to bosTau9 genome and annotated with RefSeq ARS-UCD.1.2. Quality control of the data was performed using the quality control output file from the preprocessing pipeline. Samples with low total reads, mapped reads and spike-in reads were removed. The raw count matrix (13,103 genes) was then processed with a fluctuation test to eliminate the genes which do not fluctuate more than the RNA ERCC spike-ins (i.e., only technical variation), eliminating 1,898 genes (Supplementary table 1). Fluctuated raw count matrix was used for the downstream analysis. The R package, Seurat (v4.4.0) was used to perform unsupervised sample clustering with PCA and UMAP plots. Differential expression analysis was done using the DESeq2 package (v 1.40.2) and RNA ERCC spike-ins were utilized for the data normalization; genes with less than 10 counts per sample were removed and 9,588 genes were left for the downstream analysis. For the gene to be significantly differentially expressed, FDR of 5% was used. Ggplot2 package (v 3.5.0) was used for the graphical visualization of the results. ClusterProfiler (v 4.8.3), AnnotationDbi (v 1.62.2) and bovine genome database package (org.Bt.eg.db, v 3.17.0) were used for the Kyoto Encyclopedia of Genes and Genomes (KEGG) and gene ontology (GO) analysis. Nonparametric Scheirer-Ray-Hare Test was used to perform statistical analysis of cleavage and blastocyst formation rate, followed by Dunn test. Statistical significance was considered if p< 0.05.

### Data and code availability

The raw data (BCL files) are available at Zenodo: XXXX and FASTQ files are available in EMBL-EBI BioStudies with accession number XXXXX.

## Results

### Blastocyst formation rate lowest in the ultra-low O_2_ condition

To study the effect of O_2_ concentration during *in vitro* culture of embryos, we performed 3 separate experiments. First, we performed IVM, IVF and 8 days of IVC under atmospheric 20% O_2_, hereinafter referred to as normoxia. In the second condition, IVM, IVF and 8 days of IVC were done under physiological 6% O_2_, referred to as hypoxia condition. Finally, to test the hypothesis of sequential low and ultra-low O_2_ culture, IVM, IVF and first 3 days of IVC were done under 6% O_2_ (hypoxia), after which the concentration of O_2_ in the incubator was lowered to 2%; this condition is called ultrahypoxia. Three embryos per developmental stage (zygote, 4-, 8-, 16-cell and blastocyst) were collected for the preparation of STRT-N RNA sequencing library (Figure 1A).

**Figure 1.**
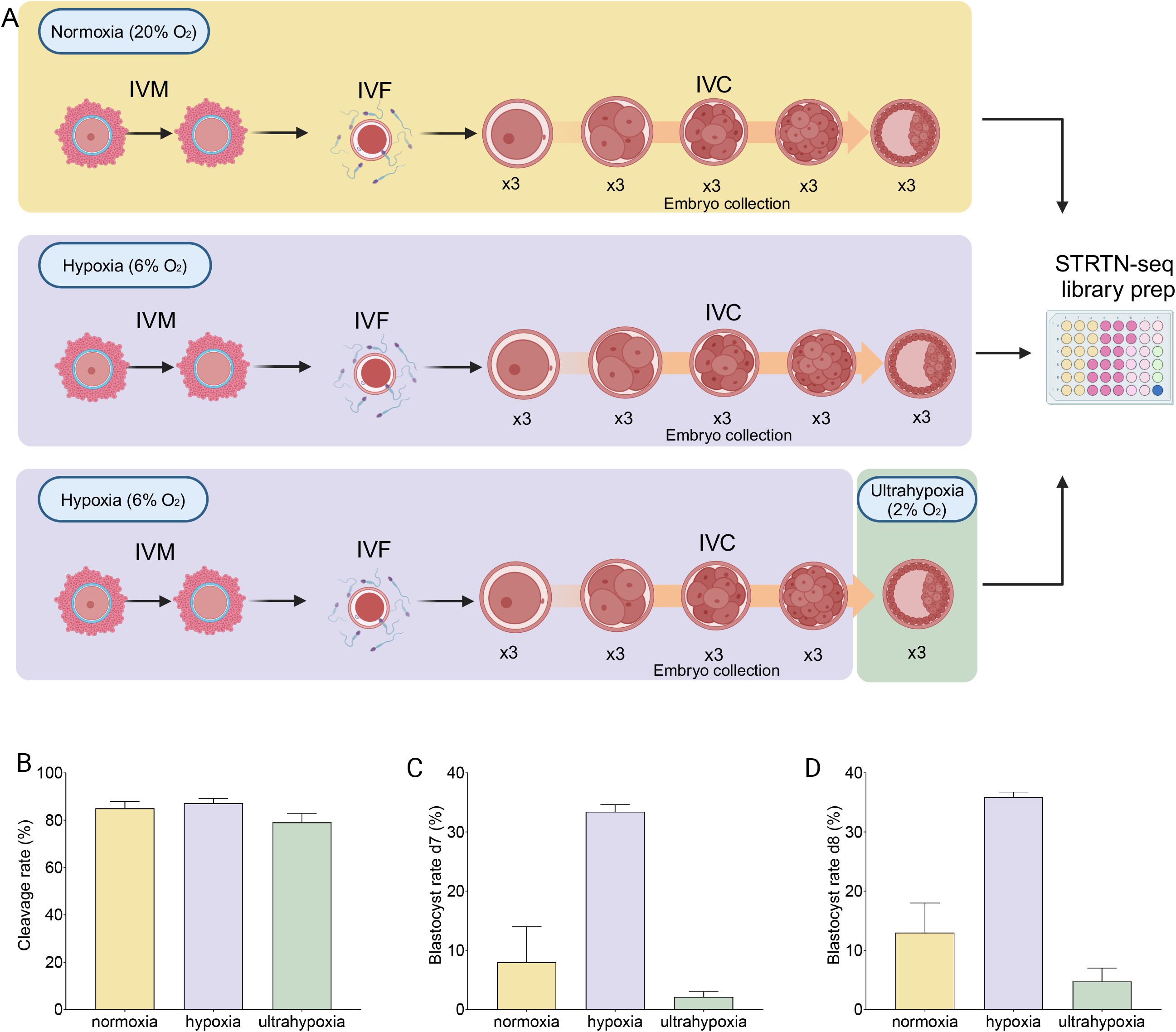
Study design and laboratory parameters. **A)** Three different experimental conditions were tested. In normoxia IVM, IVF and IVC were performed at 20% O_2_. Three embryos per developmental stages: zygotes, 4-cell, 8-cell, 16-cell and blastocysts were collected and placed in a 48-well plate containing cell lysis buffer for STRT-N seq. Hypoxia condition had the same experimental design, but O_2_ concentration used was 6%. Third condition was done by performing IVM, IVF and IVC until 16-cell stage at hypoxia 6% and then the O_2_ concentration was lowered to 2% O_2_ until blastocyst stage. Embryo collection in this experiment was done both during hypoxia stages, and blastocysts for ultrahypoxia were collected and placed in the same STRT-N seq cell lysis plate. **B)** Cleavage rate of all embryos at all conditions was calculated 24 h after the zygotes were placed in the IVC. **C)** Blastocyst formation rate on day 7 of the embryo culture was calculated including the expanded and hatched blastocysts. **D)** Blastocyst formation rate on day 8 of the embryo culture was calculated including the expanded and hatched blastocysts. Blastocyst formation rates were calculated based on the number of presumptive zygotes. Error bars represent +/-SEM.

In each experiment, a 4-well control dish was used to calculate the cleavage and blastocyst formation rate (Supplementary table 2A, B, C). Cleavage rate was determined 24 h after the embryos were placed in the culture and it was 85% in normoxia, 86% in hypoxia and 78.9% in ultrahypoxia experiment (Figure 1B). We did not observe any significant difference between the cleavage rate across conditions. In concordance with the literature, normoxia condition had a lower blastocyst formation rate both on day 7 and day 8; 8% and 13%, respectively, in comparison to the hypoxia blastocyst formation rate at day 7 and day 8 of 33% and 36%, respectively (Figure 1C, 1D). The observed difference was suggestive, but significance suffered from the low number of replicates for normoxia (n=2) (p= 0.056). Opposite to the recent reports in human and mouse (Yang *et al*., 2016; De Munck *et al*., 2019; Kaskar *et al*., 2019), we found the lowest blastocyst formation rate with 2.1% of cleaved embryos reaching the blastocyst stage at day 7 and 4.6% at day 8 in ultrahypoxia condition (Figure 1C, 1D). In comparison to hypoxia, we found that blastocyst formation rate in ultrahypoxia was statistically significant (p<0.01).

### Expression pattern of samples coincide with the development stages

The STRT-N RNA-seq is a 5’ end targeted RNA sequencing method, optimized for embryological studies. STRT-N allows distinction between degrading maternal and the newly transcribed embryonic transcripts. After sequencing, the raw data was preprocessed, and we performed quality control of the samples (Supplementary Figure 1). Due to the low number of total reads, mapped reads and spike-in reads, we removed from further analysis 3 samples: 13 (16-cell hypoxia), 26 (zygote ultrahypoxia) and 38 (4-cell ultrahypoxia) as failed reactions. To confirm the quality of the RNA sequencing, we analyzed the expression values of the housekeeping genes (*ACTB, ACTG1, ACTG2, TUBA1B, TUBA1C*) (Bilodeau-Goeseels and Schultz, 1997; Ross *et al*., 2010) and known developmental genes (*NANOG, POU5F1, GDF9, SLC34A2*) (Figure 2A). All the housekeeping genes had the expression levels trends expected in each embryo stage in accordance with the previous literature findings. *NANOG* and *POU5F1* (*OCT4*) are pluripotency markers which expression increases during the development (Canizo *et al*., 2019), and our data showed these having higher expression in blastocyst stage. *GDF9* is an oocyte secreted factor, and its expression was high in oocytes and in the early stages of the development, while it was lower after EGA in 16-cell and blastocyst stages. *SLC34A2* is an EGA marker expressed at the 16-cell stage in bovine, as also confirmed after 16-cell of development in our data. We then performed unsupervised clustering of the samples. As seen on the UMAP, early stages of development from oocytes to 8-cell stage embryos clustered together, while later stages (16-cell and blastocysts) clustered closer together and separate from the earlier stages (Figure 2B). Similarly, in the principal component analysis, the largest variance between samples (PC1) also separated the earlier stages from later stages, while PC2 separated 16-cell from the blastocysts (Supplementary Figure 2).

**Figure 2.**
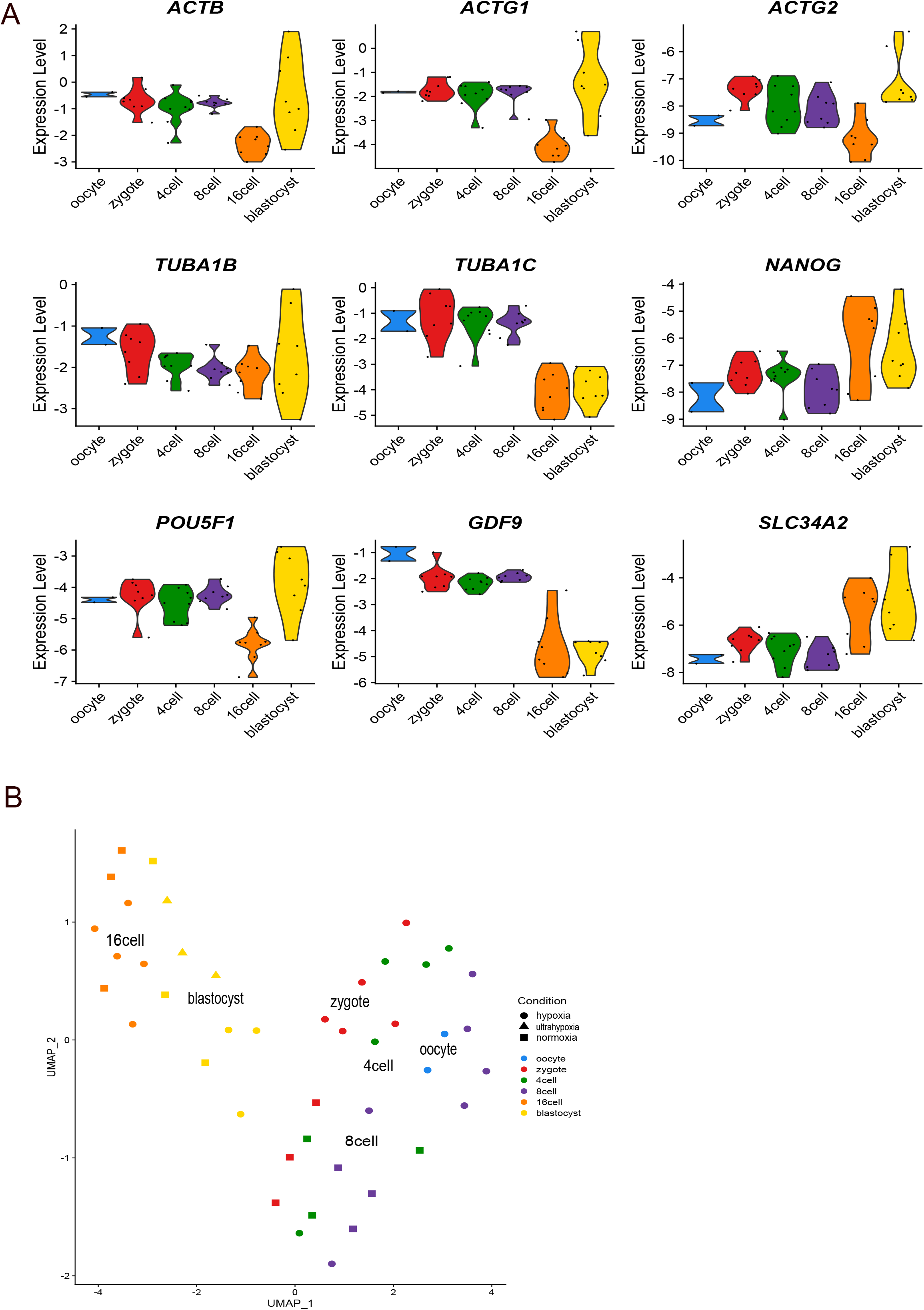
Quality control of RNA seq library and unsupervised sample clustering. **A**) Violin plots for gene expression of selected genes across different embryo development stages. Housekeeping genes *ACTB, ACTG1, ACTG2, TUBA1B* and *TUBA1C* show the expected variations in the gene expression across the development stages. The developmentally important genes *NANOG, POU5F1, GFD9* and *SLC34A* were selected for gene expression trends in this RNA sequencing library. Y-axis presents log-normalized expression levels. **B)** Unsupervised clustering was performed with R Seurat package. The UMAP plot shows that the earlier stages of embryo development cluster together and separately from the later stages: 16-cell stage and blastocyst embryos.

### Embryonic genome activation in bovine in hypoxia and normoxia

During maternal to embryonic switch, mRNA molecules from the oocyte are degraded. We did not observe any differentially expressed genes (DEG) between the zygote stage and 4-cell embryos in any conditions, different from Graf *et al*., (2014). We believe that this is due to the different sequencing and normalization methods used.

The first downregulation of genes was observed at the 8-cell stage, during which 732 genes were downregulated in the hypoxia condition, while in normoxia, only 47 genes were downregulated (Figure 3). The difference between the groups is also seen in UMAP where the normoxia 8-cell embryos cluster away from the 8-cell hypoxia embryos (Figure 2B).

**Figure 3.**
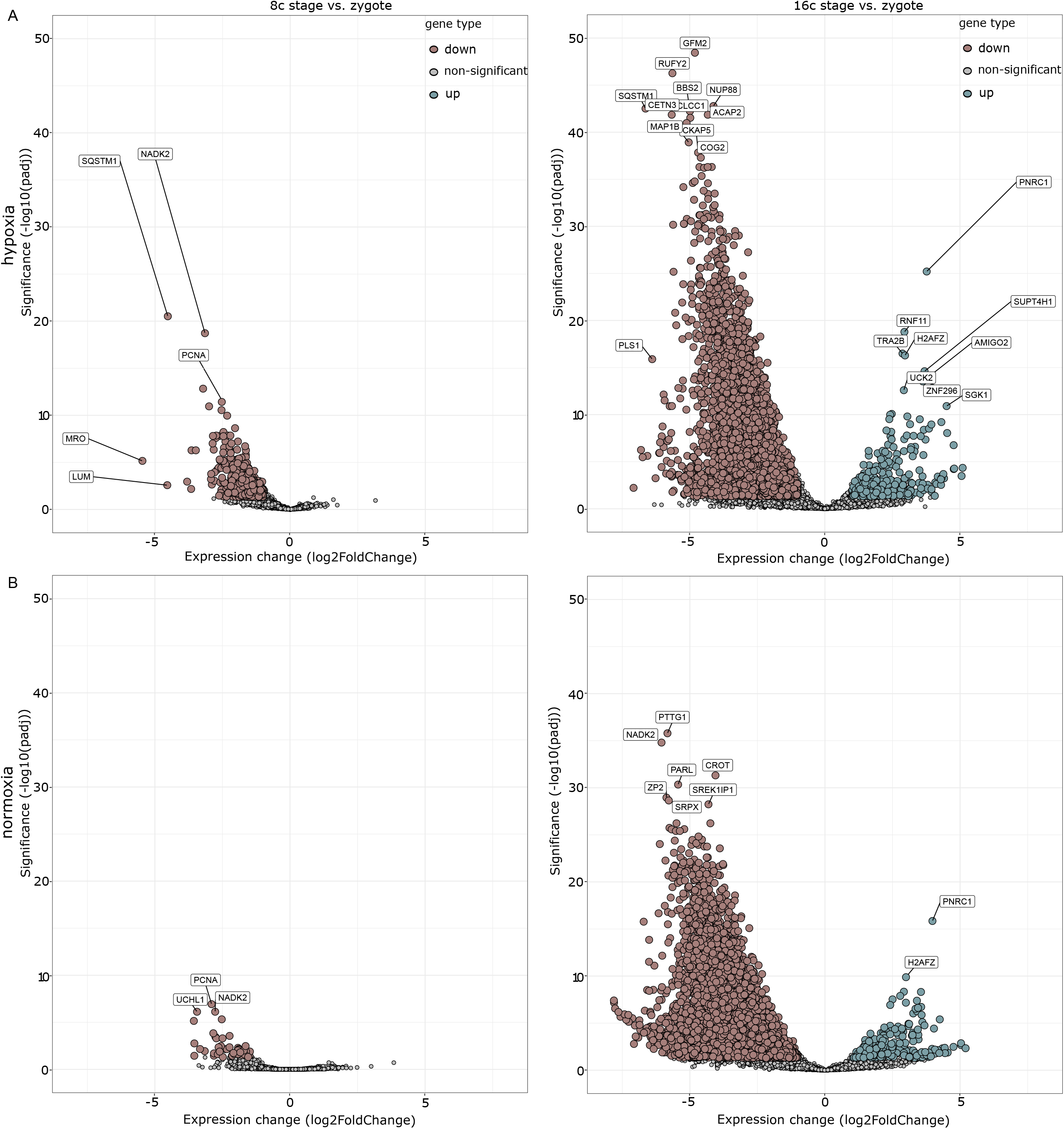
Visual demonstration of the embryonic genome activation (EGA) in bovine embryos. **A)** The track shows the transcriptional changes of embryos grown under hypoxia. The top left graph shows the differences between 8-cell stage embryos and zygotes demonstrating the first downregulation of the maternal genes (brown). On the right, we see the comparison of 16-cell embryos to zygotes, and we can see the major EGA wave in bovine, where the majority of genes is downregulated (brown) and first upregulation of the newly formed embryo transcripts appears (green). Nonsignificant changes are shown in grey. **B)** The track shows the transcriptional changes of embryos grown under normoxia. Bottom left graph presents comparison of 8-cell stage embryos against zygotes, with first downregulation of genes (brown), while the right graph demonstrates the transcriptomic changes in 16-cell stage embryos against zygotes. Downregulated genes are in brown, nonsignificant in grey, and upregulated genes in green. For the genes to be significantly downregulated or upregulated, the FDR of 5% was used.

The biggest change in DEG profile was observed between 16-cell and zygote. Here, we observed the major wave of EGA at the 16-cell stage both in hypoxia and normoxia, where also the largest maternal mRNA downregulation occurred (Figure 3).

### Gene expression during EGA

In human IVF, around 45-50% of embryos stall around 8-cell stage (Orvieto *et al*., 2022), which corresponds to the initiation of EGA in humans. We next asked if the O_2_ concentration impacts the EGA in bovine embryos. On the first look, embryos at both conditions managed to achieve EGA at 16-cell but we observed major differences in genes that were up- and downregulated at this stage in comparison to zygotes. There were 253 and 260 upregulated genes in hypoxia and normoxia, respectively. Of these 167 genes were common, while 86 and 93 were unique to hypoxia and normoxia, respectively (Figure 4A; Supplementary Table 3. Gene ontology (GO) analysis revealed that the upregulated genes in hypoxia were involved in binding, translation activity and RNA biosynthetic process and metabolism of RNA, indicating that embryos were preparing for the novel transcription, DNA replication and progress towards next stages of development (Figure 4C). In normoxia, similar GO results were seen (Figure 4E) but interestingly, in hypoxia the number of genes involved in these processes was much higher than in normoxia.

**Figure 4.**
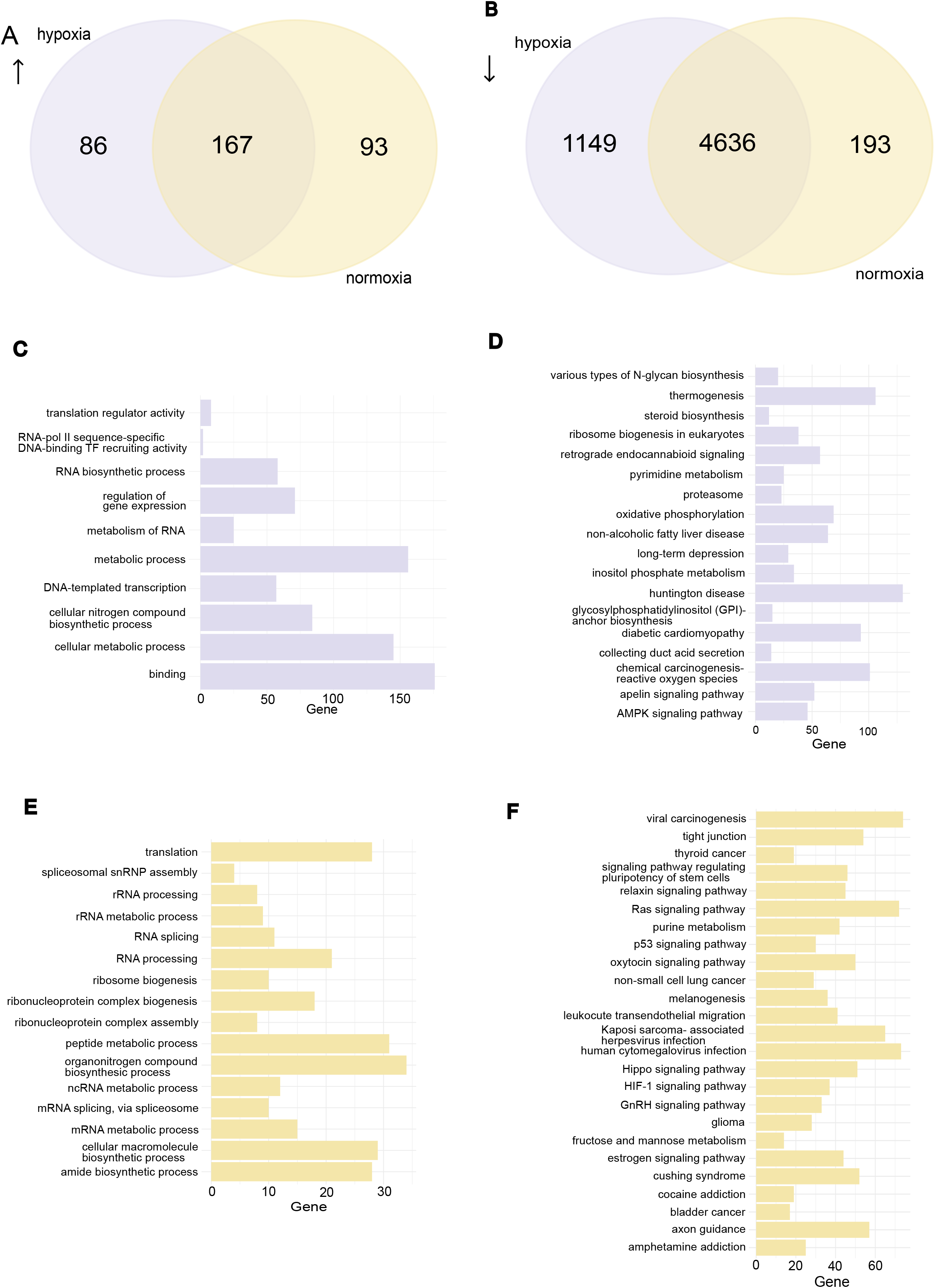
Differences in the transcription profile of 16-cell embryos grown under hypoxia and normoxia. **A)** Venn-Diagram of the upregulated genes at the 16-cell stage embryos against zygotes in hypoxia and normoxia. **B)** Venn-Diagram of the downregulated genes at the 16-cell stage embryos against zygotes in hypoxia and normoxia. **C)** Gene Ontology (GO) analysis of the genes upregulated at the 16-cell stage in hypoxia. The GO analysis includes molecular function (MF), biological processes (BP) and reactome (REAC). X-axis shows the number of genes that are identified under the GO term. **D)** Kyoto Encyclopedia of Genes and Genomes (KEGG) analysis of the downregulated genes at the 16-cell stage in hypoxia. **E)** GO analysis of the genes upregulated at the 16-cell stage in normoxia. This GO analysis includes MF, BP and REAC features. X-axis shows the number of genes that are identified under the GO term. **F)** KEGG analysis of the downregulated genes in normoxia. For term and pathway in GO and KEGG analysis, a significance threshold of 5% FDR was used.

The largest downregulation of genes appeared at the 16-cell stage where 5,785 genes or 61% of all genes in zygote stage were downregulated in hypoxia and 4,829 or 50% of genes were downregulated in normoxia (Figure 4B). To understand the function of these genes, we performed KEGG pathway analysis. In both conditions, genes related to oocyte maturation, cell cycle, autophagy and mitophagy pathways were downregulated. Surprisingly, pyruvate metabolism, carbon metabolism, 2-oxocarboxylic acid metabolism and the TCA cycle were downregulated in both conditions (Supplementary Table 4A, 4B, Supplementary figure 3, 4) which was unexpected as it is thought that pyruvate is the main energy source in the early stages of embryo development until blastocyst stage. However, in hypoxia we can also see that AMPK pathway was downregulated, suggesting that the AMP/ATP ratio inside the embryo was at a satisfactory level and that these cells do not lack energy (Figure 4D).

Besides failing to downregulate many genes in comparison to hypoxia, normoxia embryos also downregulated some of the pathways necessary for genomic integrity. The p53 signaling pathways play vital roles in genome stability, apoptosis induction and cell cycle regulation. Their downregulation in normoxia suggested that embryos grown in normoxia were deficient in the system to respond to possible DNA damage. Additionally, the HIF-1 signaling pathway was also downregulated in normoxia (Figure 4F). HIF-1 has been shown to play a vital role in EGA and metabolic adaptation as well as in expression of genes involved in angiogenesis, such as upregulation of the VEGF and GLUT-3 (Ma *et al*., 2017). The Hippo signaling pathway, which plays an important role in cell fate differentiation and cell position and is important for blastocyst formation and implantation was also downregulated in the normoxia embryos.

### Ultrahypoxia rescued the embryo energy metabolism in the blastocysts while normoxia altered it

The use of normoxia or hypoxia in the embryo culture has been long debated with recent discussions whether the best condition for the embryo culture is a sequential ultra-low O_2_ concentration. To test this hypothesis, we decided to lower the concentration of O_2_ after 16-cell stage until the blastocyst stage. This gave us the opportunity to get a transcriptional profile of all three O_2_ concentrations. We compared blastocysts in each condition against the 16-cell stage embryos. A major upregulation of genes was seen both in hypoxia (1,991) and ultrahypoxia (1,967) blastocysts while in normoxia blastocysts upregulation of only one quarter of those (539) was observed (Figure 5A-D). Interestingly, in terms of transcriptional downregulation the largest difference was seen in ultrahypoxia (1,338) in contrast to normoxia with 663 genes, and only 222 in hypoxia (Figure 5D).

**Figure 5.**
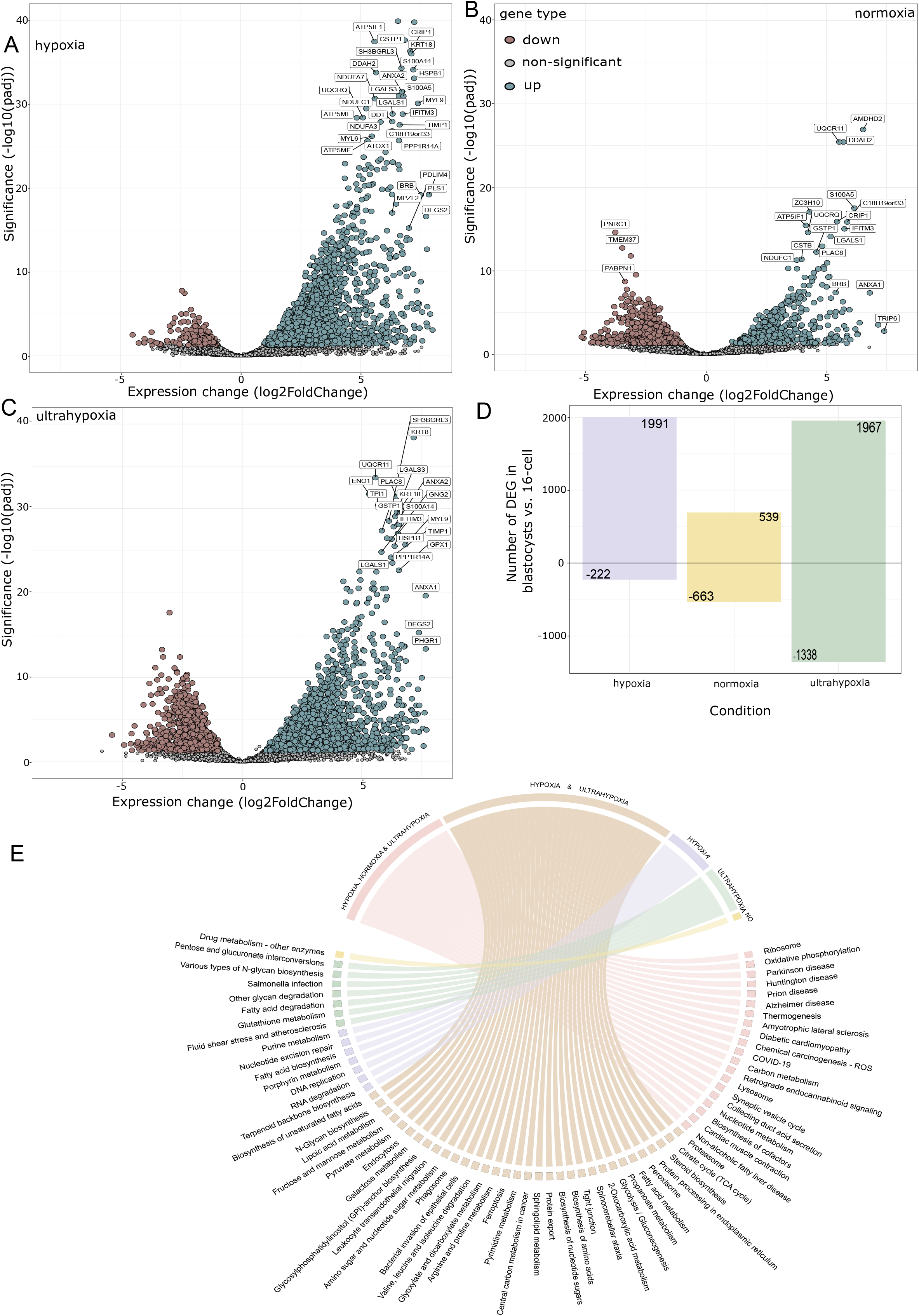
Comparison of blastocysts transcriptomic profiles grown under hypoxia, normoxia and ultrahypoxia. **A)** Volcano plot representation of the transcriptional difference in the blastocysts grown under hypoxia against 16-cell stage hypoxia embryos. **B)** Volcano plot representation of the transcriptional difference in the blastocysts grown under normoxia against 16-cell stage normoxia embryos. **C)** Volcano plot representation of the transcriptional difference in the blastocysts grown under ultrahypoxia against 16-cell stage hypoxia embryos. Downregulated genes are shown in brown, not significant in grey and upregulated genes in green. **D)** Number of differentially expressed genes (DEG) in blastocysts against 16-cell embryos across the 3 experimental conditions. **E)** Chord diagram of the KEGG pathways upregulated at the blastocysts stage against 16-cell stage embryos in all 3 conditions, showing which KEGG pathways are shared between all 3 conditions (pink), only in hypoxia and ultrahypoxia (beige), only in hypoxia (purple), only in ultrahypoxia (green) and only in normoxia (yellow) marked as NO on the figure.

Next, we wanted to define the influence that O_2_ concentration had on the genes regulating energy metabolism of the blastocysts. At this time of development, blastocysts should be powered mostly by glucose, making glycolysis the key energy pathway. We observed that all 3 groups had genes upregulated in the oxidative phosphorylation pathway, but only hypoxia and ultrahypoxia blastocysts had pyruvate metabolism and the TCA cycle genes upregulated (Figure 5E). The largest difference in the metabolism related gene expression could be seen in the glycolysis/gluconeogenesis pathway that was only active in hypoxia and ultrahypoxia. In addition, genes related to amino acid biosynthesis of arginine, proline, valine, leucine and isoleucine were present in both hypoxia and ultrahypoxia while absent from the normoxia condition. Genes involved in lipid metabolism including fatty acid biosynthesis, biosynthesis of unsaturated fatty acids, steroid biosynthesis and sphingolipid metabolism were also only present in the hypoxia and ultrahypoxia conditions (Figure 5E). Overall, even though the blastocyst formation rate in our experiment was the lowest in the ultrahypoxia condition, the transcriptomic profile of the blastocysts grown in the ultrahypoxia was similar to the hypoxia condition with the key genes in energy metabolism necessary for the further cell differentiation and later organogenesis upregulated in ultrahypoxia. Normoxia blastocysts showed that they were much slower to achieve the full transcriptional awakening at the blastocyst stage and had altered expression of energy metabolism genes that can have further consequences for the development.

## Discussion

The oxygen level used for *in vitro* embryo culture has been a topic of discussions for many years. The main question is whether culturing embryos under physiological O_2_ levels (hypoxia) is beneficial in comparison to culturing embryos under atmospheric O_2_ (normoxia). There is an increasing amount of evidence suggesting that hypoxia is superior compared to normoxia. In human normoxia, IVF success rate of embryo cultures is around 30%, while hypoxia increases the success rate up to 32-43% (Bontekoe *et al*., 2012). Kovačič and Vlaisavljević (2008) showed that the proportion of good quality embryos at day 3 and higher blastocyst formation rate were achieved in hypoxia. Van Montfoort *et al*. (2020) included both fresh and frozen embryos (5-7 years follow up) in their study and found hypoxia yielding significantly more good quality embryos and live births in comparison to the normoxia group using day 2-3 embryos. On the contrary, De Los Santos *et al*., (2013) did not find any difference in the clinical pregnancy rate between the two conditions using day 2-3 embryos. This is possibly due to different ways of calculating the pregnancy rate and no information was provided on the period of the follow up with the study participants and if more embryos were left cryopreserved.

Due to ethical limitations to use human embryos, literature lacks evidence about the impact O_2_ concentration has on the transcriptomic level. To reach molecular understanding, mouse embryos have been used. Kelley and Gardner (2019) tested the effect of normoxia and hypoxia and found that normoxia decreased the blastocyst formation rate, but also altered the energy metabolism by decreasing the uptake of amino acids leucine, methionine, and threonine, as well as the uptake of glucose and aspartate. Belli *et al*., (2019) tested the impact of normoxia on mitochondria. They showed that blastocysts derived from normoxia had a lower mitochondrial DNA copy number and fewer mitochondria compared to the hypoxia.

In this study, we chose bovine embryos as a model organism due to their closer resemblance to humans. We employed STRT-N RNA-seq to differentiate between maternal and embryonic transcripts. Proper data normalization is crucial in embryology studies, as most methods assume that the majority of co-expressing genes are not differentially expressed between samples. However, this assumption does not hold during embryo development because of significant transcriptional shifts between stages by maternal transcript degradation (Katayama *et al*., 2013). By using the appropriate normalization technique, specifically the ERCC Spike-ins, we successfully accounted for sample heterogeneity in our data.

We showed that normoxia slows down the degradation of the maternal transcripts during the EGA. Degradation of the maternal transcripts and switch to the embryonic transcription are prerequisites to normal development. Normoxia influenced EGA in bovine embryos by failing to downregulate 1,149 genes at 16-cell stage. Moreover, we observed downregulation of the HIF-1 signaling pathway that has been highlighted as the cellular response to hypoxia, but, interestingly, HIF-1 also regulates transcription of many genes involved in glucose uptake, glycolysis, angiogenesis, proliferation, and autophagy (Bracken *et al*., 2003). The influence of the HIF-1 signaling pathway on the development has been studied in mouse and bovine embryos. In mouse, it has been shown that HIF-1 has a direct upregulating impact on the expression of several major EGA genes (Yao *et al*., 2024) and that HIF-1 is degraded in normoxia, confirming the previous reports of HIF-1 degradation by O_2_ concentration (Salceda and Caro, 1997; Lando *et al*., 2002; Marxsen *et al*., 2004). One of the HIF-1 targets is vascular endothelial growth factor (VEGF) that is important for angiogenesis and the connection of the embryo and endometrium during embryo implantation, and it has been shown to have an impact in the recurrent implantation failure (Taghizadeh *et al*., 2022). Downregulation of HIF-1 signaling in normoxia embryos can have a detrimental effect on the development potential of embryos, although its role may not be evident in the first few days of development.

In our study, the Hippo pathway and signaling pathways regulating stem cell pluripotency were also downregulated at the 16-cell stage in normoxia embryos. The Hippo pathway works through the two activators, YAP and TAZ, and has been shown to regulate cell differentiation and specialization of trophectoderm (TE) and inner cell mass (ICM) (Yildirim *et al*., 2021). Large tumor suppressor kinase (LATS) is known as Hippo pathway inhibitor. In mouse, suppressing LATS1/2 expression during preimplantation development by siRNA injection disrupts Hippo signaling pathway, leading to cell fate misspecification in ICM and ultimately the failure of embryos to undergo gastrulation (Lorthongpanich *et al*., 2013).

We also observed that normoxia disrupted the expression of genes involved in energy metabolism of the blastocysts. While blastocysts’ main energy source should be glucose, we found that in normoxia blastocysts genes involved in glycolysis, gluconeogenesis, and synthesis of amino acids were not significantly upregulated. The lack of energy could partially explain lower developmental competence and, for example, lower cell counts in blastocysts observed in several animal studies (Belli *et al*., 2019). This question becomes even more relevant considering the new findings that glucose gradient might be involved in gastrulation and mesoderm establishment (Bulusu *et al*., 2017; Cao *et al*., 2024).

In addition to testing the difference between hypoxia and normoxia, we tested the effect of sequential ultralow O_2_ concentration on the bovine embryo development. We compared the transcriptional profile of the ultrahypoxia blastocysts and hypoxia blastocysts. We found a larger number of downregulated genes in the ultrahypoxia blastocysts when compared against 16-cell stage than in the same comparison done with hypoxia blastocysts. However, the energy metabolism profile, as estimated by upregulated DEGs, of the ultrahypoxia blastocysts was similar to the hypoxia. Contrary to normoxia blastocysts, ultrahypoxia blastocysts had upregulated genes related to glycolysis, amino acid synthesis and fatty acids biosynthesis. Morin *et al*. (2018) performed RNA sequencing on ICM and TE from aneuploid human blastocysts. They did not find differences in the transcription profiles of the blastocysts grown in monophasic 5% or sequential 5-2% O_2_. Yang *et al*. (2016) compared the 2%, 5% and 20% O_2_ levels in human embryo development. They also investigated the expression of the genes related to the TE and placentation and showed the difference between the 20% O_2_ and hypoxia conditions but saw no difference between 5% and 2% O_2_ groups.

De Munck *et al*. (2019) performed a randomized trial where human embryos on day 3 of development were either continuously grown under 5% O_2_ or moved to 2% O_2_. They found no difference in the blastocyst formation rate and proportion of good quality embryos. Yang *et al*. (2016) did not observe any difference in the blastocyst formation rate between 5% and 2% O_2_, but they did find lower cell number in the blastocysts grown at 2%. In our experiments with bovine embryos, the lowest blastocyst formation rate was in the ultrahypoxia condition, but in the current study the cell number in the blastocysts across the conditions was not counted.

Despite the similar transcriptomic profiles between ultrahypoxia and hypoxia blastocysts observed in our study and in previous human reports, some studies call for caution of switching to the 2% O_2_. Feil *et al*. (2006) tested these three O_2_ conditions in the mouse embryo development and showed that fetal weight on day 18 was lower in embryos cultured at 2% O_2_. Similarly, Kaser *et al*. (2016) showed some adverse effects of 2% O_2_, as they observed fewer cells in the blastocysts as well as in the absorption of amino acids and their abundance and expression of MUC1 in the TE.

In summary, we provided transcriptional profiles of bovine embryos grown in normoxia, hypoxia and hypoxia/ultrahypoxia. Normoxia had a detrimental influence on the transcriptomic switch of the embryo by slowing down the downregulation of the maternal transcripts during EGA, but also by downregulating developmentally critical pathways and failing to upregulate proper energy metabolism. We hypothesize that all of these might be reasons for the normoxia grown embryos resulting in lower number of live births in humans. Ultrahypoxia blastocysts had similar profiles to hypoxia by upregulating the energy metabolism needed to fuel the blastocysts. Despite this, we saw the lowest blastocyst formation rate in ultrahypoxia, opposite to previous reports in human. As there is no clear benefit of lowering the O_2_ to 2% and considering the extra costs needed for such ultrahypoxia culture, we propose monophasic low O_2_ concentration as optimal for *in vitro* embryos culture systems.

## Authors’ roles

N.B. study design, embryology experiments, RNA sequencing experiments, data analysis, writing the original draft; M.I. embryology experiments, manuscript revision; G.Y., B.Y. preprocessing of RNA sequencing data; S.K., K.L., T.T., T.O., A.K., A.S., supervision, study design, review & editing of manuscript; C.D., supervision of data analysis, review & editing of manuscript; J.K., conceptualization, supervision, writing, review & editing, project administration and funding acquisition.

## Acknowledgments

We would like to thank Biomedicum Functional Genomics Unit (FuGU), University of Helsinki, for library sequencing services. The bioinformatics data analyses were performed on CSC – IT Center for Science, (Finland).

## Supplementary data description

**Supplementary Figure 1. Quality control analysis of the STRT-N RNA sequencing library**.

**A)** Graph showing total number of mapped reads. **B)** Total number of ERCC Spike-in reads per sample. **C)** Quality check of the 5’-end spike-in rate. **D)** Mapped rate calculated by dividing the total number of reads with total number of mapped reads. **E)** Mapped/Spike-in reads, a quality check of the overall amount of mRNA present in each sample. **F)** Coding 5’-end of the mRNA molecules. On the x-axis of all the samples we have developmental stages. Hypoxia is shown in red, normoxia in blue and ultrahypoxia in green.

**Supplementary Figure 2. Principal component analysis**.

Principal component analysis with all the samples used for the downstream analysis. PC1 shows the biggest variance between samples according to the developmental stages of the embryo development, separating the earlier stages of development: oocytes, zygotes, 4- and 8-cell embryos; from the later stages: 16-cell embryos and blastocysts. PC2 separates blastocysts from the earlier stages of embryo development.

**Supplementary Figure 3. TOP 40 downregulated KEGG pathways in 16-cell embryos in hypoxia**.

Top 40 KEGG pathways separated based on their p. adjusted value are shown on the y-axis. The x-axis represents the count of genes involved in the pathways, while the color of the circle indicates the significance value.

**Supplementary Figure 4. TOP 40 downregulated KEGG pathways in 16-cell embryos in normoxia**.

Top 40 KEGG pathways separated based on their p. adjusted value are shown on the y-axis. The x-axis represents the count of genes involved in the pathways, while the color of the circle indicates the significance value.

**Supplementary Table 1. Fluctuated gene count matrix**

Fluctuated raw count matrix used for the downstream analysis of differential expressed genes, GO and KEGG analysis. Fluctuated raw count matrix contains only the genes that have higher variance then the RNA ERCC spike-ins.

**Supplementary Table 2. Cleavage and blastocyst formation rate**.

Calculation of cleavage and blastocyst formation rates in normoxia, hypoxia and ultrahypoxia.

**Supplementary Table 3. Upregulated genes at the 16-cell stage embryos in hypoxia and normoxia**.

Full list of the upregulated genes at 16-cell embryos compared to the zygote stage. There are 167 common genes that are upregulated both in hypoxia and normoxia. There are 87 upregulated genes unique to hypoxia, while 93 upregulated genes are unique to normoxia.

**Supplementary Table 4A. Complete list of KEGG pathways downregulated at the 16-cell embryos in comparison to zygotes in hypoxia**.

Full list of KEGG pathways downregulated at the 16-cell stage embryos in hypoxia.

**Supplementary Table 4B. Complete list of KEGG pathways downregulated at the 16-cell embryos in comparison to zygotes in normoxia**.

Full list of KEGG pathways downregulated at the 16-cell stage embryos in normoxia.

